# Generation of biobased, biodegradable non-woven straw mats for multiple environmentally friendly uses

**DOI:** 10.1101/2024.08.07.596360

**Authors:** Jonathan Gressel, Sigal Hoffmann

**Affiliations:** HiCap Formulations (Israel) Ltd, Rehovot, Israel; Department of Plant and Environmental Sciences, Weizmann Institute of Science, Rehovot, Israel

**Keywords:** Straw, lignin, biodegradable adhesives, aquaculture, weed control, insulation

## Abstract

Two billion tons of grains straws are produced annually, most of which has a negative ecological value. A small proportion is fed to ruminants as a low calory roughage. Grain straws had been burnt to prevent pathogen spores from over-wintering, now requiring greater fungicide use if left on soil surface, or more fertilizer use when the straw is plowed under and then binds nutrients. Lignin from paper making had been dumped, but is now finding limited uses, including as a glue in plywood manufacture. We propose to find the right ratio of lignin and other biodegradable adhesives coated on straw along with ascertaining the optimal pressures and temperatures for binding the adhesives and them cross-link straw fibers into mats.

These slowly-biodegradable mats can be as: 1. Filters surrounding fish pens, binding pollutants from fish excrements, which are metabolized along with the straw by periphyton into material edible by fish as well as zooplankton eaten by fish. These filter mats may also prevent the movement of parasites into or out of the pens; 2. Mats used for erosion control on bare slopes until vegetated; 3. As insulation material in construction, where the carbon will be sequestered for decades; 4. As a compostable packaging material replacing polystyrene and bubble wrap; 5. Weed-preventing soil covering in organic and conventional agriculture. Such mats can be doped with ammonium and potassium as well as calcium and magnesium to be slowly released as fertilizer. Thus, by combining a negative environmental value waste materials with an adhesive we can generate mats that have very positive environmental benefits.

## Introduction

About half of the above ground biomass of grain crops is wasted: the straw that bore the grain. Most of the *ca*. two billion tons of the rice, wheat, and maize straw produced annually in the world has a negative economic value. Much of it had been burnt after harvest until environmental rules prohibited this custom. Straw temporarily binds mineral nutrients after being plowed-under, often requiring additional fertilizer in the following crop, with negative economic and environmental consequences. Small amounts of straw are fed to ruminants as roughage or as an extender to animal feeds, but very little caloric value is derived from it. Most of what remains in straw by harvest time are hemicelluloses (mainly xylans) and cellulose, but their enzymatic biodegradation is heavily prevented by a smaller component of lignin (Gressel, Vered et al. 1983).

The aquaculture of fish results in large scale production of fish excrements. The excrements of caged farmed fish pollute surrounding waters, and are a waste of utilizable mineral nutrients. The soluble wastes of intensive tank produced fish are filtered through a sophisticated system that utilizes bacteria on beads to denitrify ammonia and urea to dinitrogen gas (Shnel, Barak et al. 2002), a loss of the considerable energy used to produce urea and ammonia. Straws and other agricultural wastes are added to fish ponds in the Far East to absorb urea and ammonia. A biofilm of periphyton bacteria and fungi grow on the straw in fish ponds using the lignin and carbohydrates from the straw as their carbon source together with the bound ammonia, nitrates and soluble phosphorous emanating from fish excretions as sources of nitrogen, providing a complete medium for periphyton growth (Umesh, Shankar and Mohan 1999) (Mridula, Manissery et al. 2005) (Thomas, Lalramchhani et al. 2020). Fish directly consume to periphyton or indirectly via crustaceans and other zooplankton derived their nutrition from the periphyton, requiring less exogenous feed (Rai, Yi et al. 2008). There has been no cost-effective solution yet to prevent the pollution from fish penned in cages in the sea and rivers.

We suggest that it may be possible to use mat filters made of a slowly biodegradable adhesive cross-linked straw to cover the bottom and sides of fish pens that will allow water movement through the enclosures. The biofilms (periphyton) that develop on filters can slowly degrade filter material, while absorbing ammonia from the fish in penned enclosures. The filter mats with the sticky periphyton that develop on them should reduce or prevent movement of parasites and pathogens through the filters, just as rather open air-conditioner filters trap dust particles. Water would still flow through the mats allowing oxygenated water to sustain the fish.

Mats made by enclosing straw in plastic netting are already used in erosion control on slopes and as a ground cover to prevent weed growth and save water in agriculture. Such loose mats cannot be used as vertical water filters as the straw would sag, and leave plastic residues after biodegradation.

There is a vast literature on the production of biobased adhesives that are biodegradable. The market for such products is broad and large (Anonymous 2022). For at least four decades there have been publications describing adhesives based on waste lignin products, (Young, Fujita and River 1985) (Frazier 2023) (Yang, Gong et al. 2023), (Mili, Hashmi et al. 2022) along with lignin composites with glyoxal (Siahkamari, Emmanuel et al. 2022), polyurea (Liu, Fang et al. 2020), as well as epoxy (Li, Gutierrez et al. 2018), all suggesting that these adhesives should be used for gluing wood, especially in plywood production. Despite the publications, we have been able to locate only two companies actually commercializing lignin based adhesives. Many other inexpensive bioproducts have been modified to enhance their use of adhesives, especially to render them hydrophobic. These include hydrophobic starch for gluing paper (Watcharakitti, Nimnuan et al. 2023), urea-oxidized starch (with nano-titanium dioxide) (Zhao, Peng et al. 2018), polylactic acid (Ranakoti, Gangil et al. 2022), and many others recently reviewed (Nordstrom, Demrican et al. 2018, Antov, Savov and Neykov 2020). None of these articles suggest using adhesives to produce non-woven straw mats, but powdered straw has been graft polymerized with acrylamide to produce 100 mesh particles that absorb 8 times their weight in water (Wan, Huang et al. 2013).

We report below the small-scale experimental construction of non-woven, cohesive straw mats of different densities, porosities, and thicknesses made by cross-linking the straw using biodegradable adhesives. Such mats would not leave plastic residues, and the cross-linkers would delay the biodegradation of the straw polysaccharides and lignin from the mats, allowing them to perform their functions for longer periods of time.

These mats are being developed by us for the following main uses:

1. As filters in fish farming to remove fish excrements, preventing pollution and recycling the minerals and organic material back to fish feed via the periphyton, which will slowly degrade the straw and adhesive as carbon sources. Such filters may also preclude the movement of fish pathogens and parasites into/out of fish cages;
2. To replace plastic-netted straw mats in erosion control on bare slopes, and seed protection mats for lawn development.
3. For use as light impenetrable mats in agriculture to prevent weed growth and prevent soil drying, with crop seedlings transplanted through the mats. We will also evaluate whether these mats can be doped with specific minerals to provide a slow release of fertilizers, preventing pollution.

## Materials and Methods

Wheat straw was kindly provided by Dr Omri Sharon. The stage of a commercial heat press was modified to hold a 5 cm deep 38×38 cm wood frame with a sheet of polytetrafluoroethylene (PTFE) secured to the base of the frame (Fig.1). The frame can contain up to 250 g of straw treated with varying amounts of adhesive. Only those adhesives tested that cross-linked the straw into sufficiently stiff, non-friable mats that did not disintegrate in water are described below.

**Figure 1.**
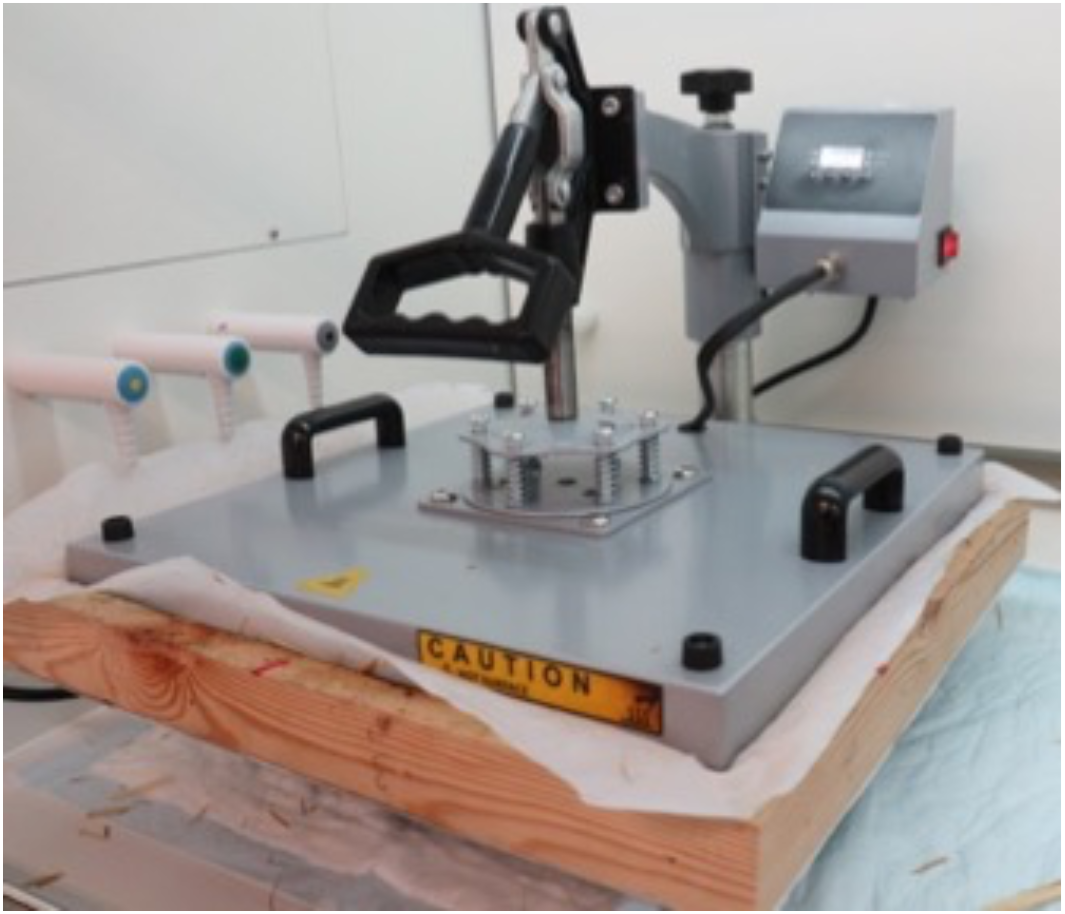
A T-shirt printer hot press modified to make experimental-sized straw mats. Parts were removed to accommodate a 38×38 cm (internal dimensions) wooden form with a Teflon sheet attached to the lower side, containing 250 g of straw spray-coated with binder and covered with a Teflon or fiberglass sheet. The lower heat plate (stage) is obscured by the wooden frame. The temperature and time pressed varied according to the binder used.

### The adhesives and their use

*LigninTack* is a water soluble proprietary adhesive composed of lignosulfonate and chitosan produced by Ribler GmbH, Plieninger Straße 5870567 Stuttgart-Möhringen, Germany. The adhesive has a solid content 9.2%.. After mixing adhesive with straw, the mixture was heat-pressed at 110C for one hour to evaporate the water and perform the binding.

*OC BioBinder*^*TM*^ *Lily 1450* is a proprietary adhesive composed of L-(+)-lactic acid, (2S)-2-hydroxypropanoic acid with 1,2 benzisothiazol and 5-chloro-2-methyl-3(2H)-isothiazolone with 2-methyl-3(2H)-isothiazolone as preservatives, produced by OrganoClick AB Linjalvägen 9 SE-187 66 Täby Sweden. The adhesive has a solid content of ca. 28%. Treated straw was subjected the heat press for 1 hour at 150°C.

*ST6515* is a proprietary adhesive based on polyurethane chemistry with 5-chloro-2-methyl-4-isothiazolin-3-one and 2-methyl-2h-isothiazol-3-one added as preservatives produced by Scitech Adhesives and Coating, Flint, Wales. The adhesive had a solid content of 40%. The mixture was subjected to 150°C heat treatment for 1H in the heat press.

*Wet strength* was crudely measured by placing the filters, weighted down, in water for 5 days.

### Prevention of weed seed germination

A plot 2m x 0.8m with a rich potting soil was seeded with 4 grams (ca. 8000) *Amaranthus palmeri* seeds. The seeds were kindly provided by WeedOut Ltd. Three 18×18 cm mats with a hole drilled in center with a keyhole drill were placed in the plots and transplant plugs of tomato (*Solanum lycopersicum*) were planted in a manner that completely filled the holes in the mats. Two tomato plants were interspersed between the mats as controls. Weed counts were performed 9 weeks after planting when the weeds began to set seed, necessitating termination..

## Results

A series of commercially available bioadhesives were tested in our system. Three examples showing positive results are shown in Fig. 2.

**Figure 2.**
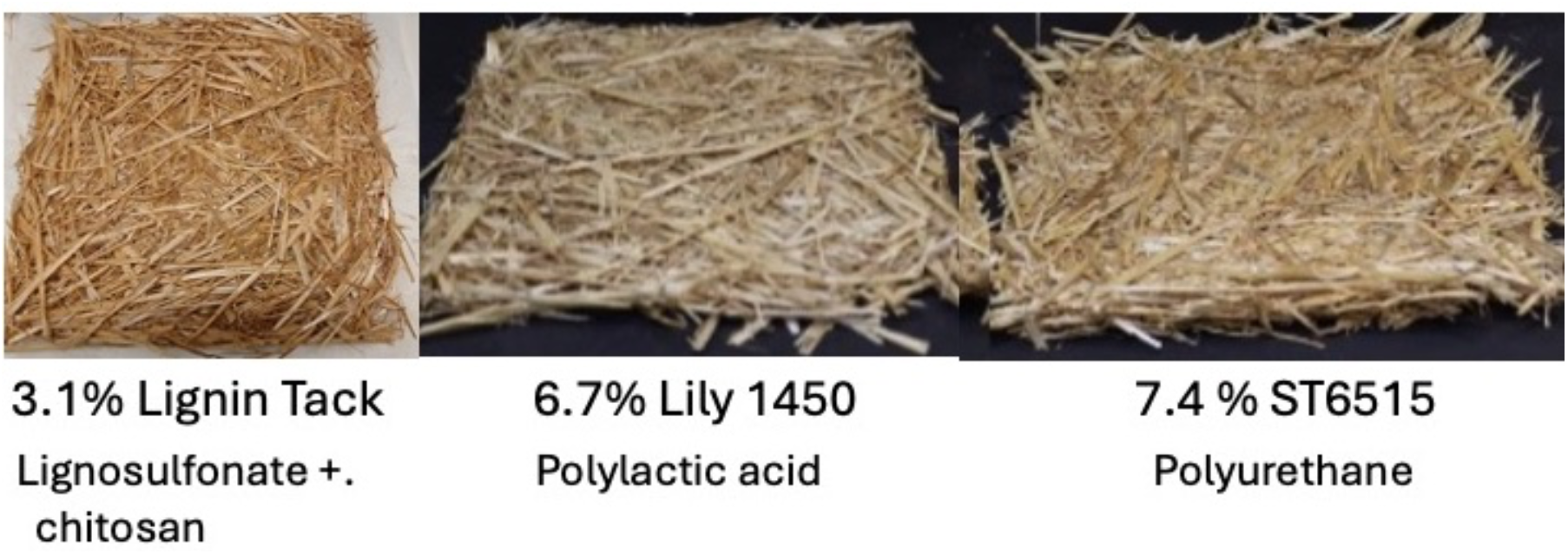
Examples of straw mats produced with different adhesives.

Two concentrations 17%w/w and 31%w/w of the *OC BioBinder*^*TM*^ *Lily 1450* polylactic acid based adhesive showed sufficient stiffness for a variety of uses including biofilters for placing on fish cages as did the 7.4%w/w ST6515 polyurethane cross-linked straw. The low 5.1% w/w Lignin-Tack adhesive provided a looser binding, for which there are many potential uses such as soil cover. Higher concentrations provide greater stiffness.

Poured water readily ran through the mats showing that they are not an impediment to water movement and thus they have a potential as filters.

The filters maintained their structure when submerged in water for 5 days (Fig. 3).

**Figure 3.**
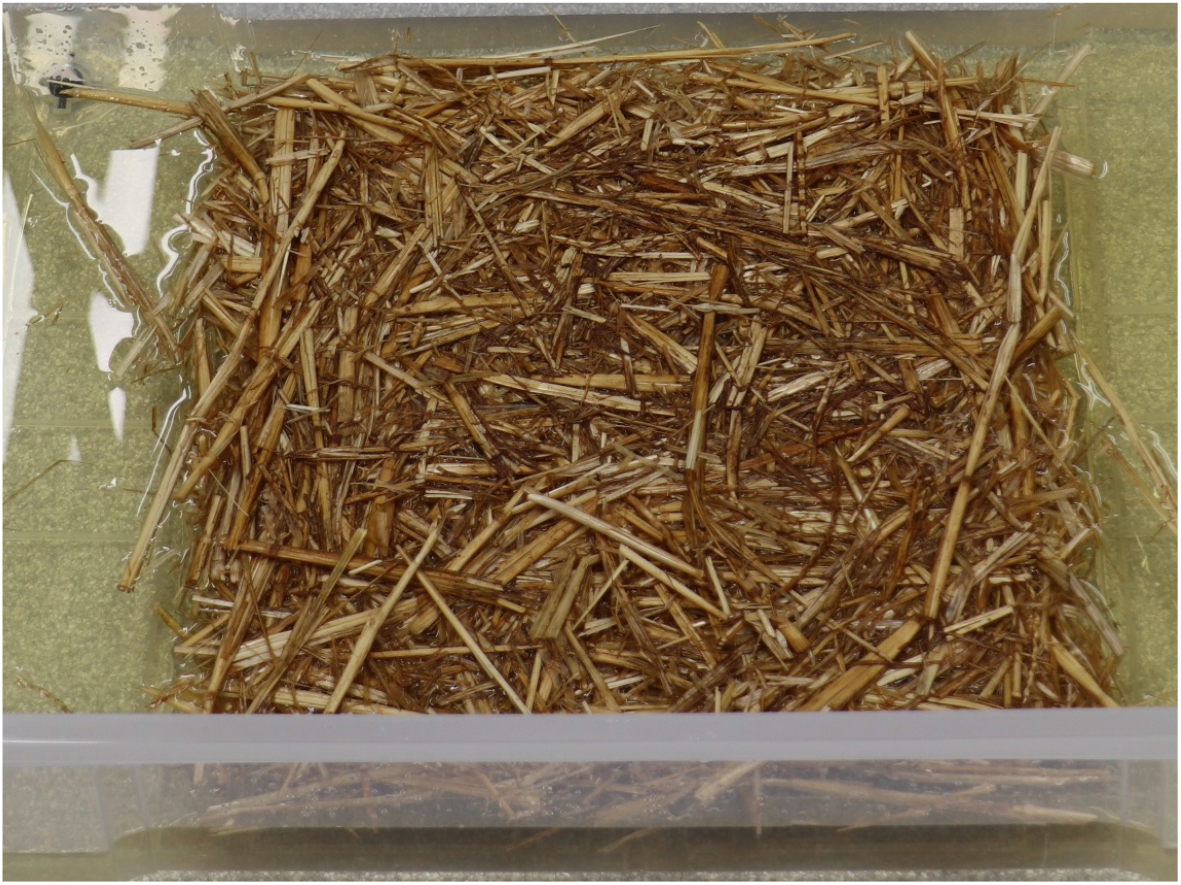
Example of a filter maintaining most of its structure when soaked in water – in this case a filter made using 13% w/w *OC BioBinder*^*TM*^ *Lily 1450* polylactic acid based adhesive is shown. They did not disintegrate when removed from the water – the crosslinking did not dissolve.

### Prevention of weed seed germination

The experiment at the time of planting is shown in Fig. 4A. The *Amaranthus* seeds germinated within 5 days as a dense mat, followed by considerable self thinning. At the time of terminating the experiment the *<u>Amaranthus</u>* pants were about 50 cm tall, many with the thick stems the prevent mechanical harvesting. Throughout the experiment no *Amaranthus* seedling penetrated the mats (Fig 4B), demonstrating their potential utility.

**Fig. 4.**
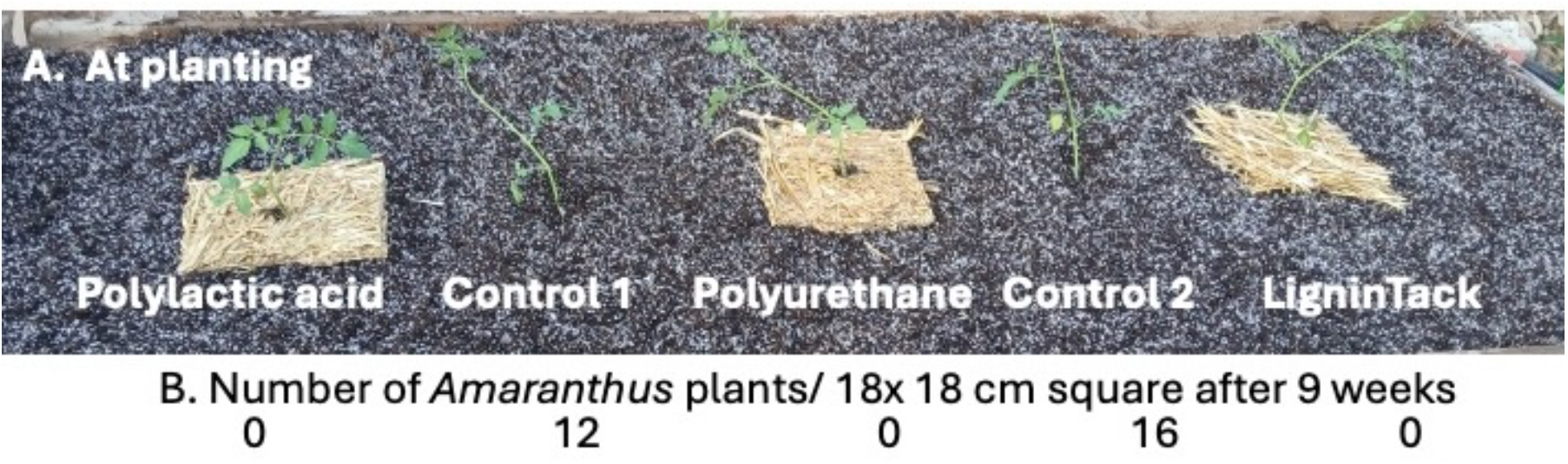
Straw mats prevent emergence of the pernicious weed *Amaranthus palmeri*. A. the experimental set-up at time of planting; B. Weed seedling count after 9 weeks.

## Discussion

It is fascinating that there is no prior indication in the scientific literature of the production of non-woven straw mats using an adhesive – whether biobased or synthesized from hydrocarbons. The production of such straw mats should be subject to receiving carbon credits, both due to reduced use of fossil fuels and the delayed release of carbon dioxide due to the slower degradation of the straw. As discussed in the introduction, it is well documented that periphyton growing on straw reduces the amount of feed required for aquaculture. Farmed fish are the most feed efficient form of animal protein and future food security requires that they should a larger proportion of the animal protein in the human diet. The use of straw accentuates thus reasoning as it releases more feedstocks for other purposes.

Straw matting for erosion and weed control reduces run-off, promotes water retention by shading the soil and reduces the need for using herbicides. If the straw is doped with fertilizers it can release the fertilizer slowly, as needed by the crop, reducing the amount percolating into the underground water table or running off into surface water. When the mats are eventually incorporated into the soil the residues will enrich the soil organic matter with humic compounds.

Thus, bio-adhesive cross-linked straw mats can make a major contribution to the green bio-economy.

## Acknowledgements

Some aspects of this research were supported by a research grant from the Yotam project and the Weizmann institute sustainability and energy research initiative”, and by “a research grant from the Tom and Mary Beck Center for Advanced and Intelligent Materials at the Weizmann Institute of Science, Rehovot, Israel”. We are grateful to Dr Filipe Natalio for graciously providing laboratory space and suggestions regarding this research.

